# Predicting behavior from eye movement and whisking asymmetry

**DOI:** 10.1101/2021.02.11.430785

**Authors:** Ronny Bergmann, Keisuke Sehara, Sina E. Dominiak, Jens Kremkow, Matthew E. Larkum, Robert N.S. Sachdev

## Abstract

Navigation through complex environments requires motor planning, motor preparation and the coordination between multiple sensory–motor modalities. For example, the stepping motion when we walk is coordinated with motion of the torso, arms, head and eyes. In rodents, movement of the animal through the environment is often coordinated with whisking. Here we trained head fixed mice – navigating a floating Airtrack plus maze – to overcome their directional preference and use cues indicating the direction of movement expected in each trial. Once cued, mice had to move backward out of a lane, then turn in the correct direction, and enter a new lane. In this simple paradigm, as mice begin to move backward, they position their whiskers asymmetrically: whiskers on one side of the face protract, and on the other side they retract. This asymmetry reflected the turn direction. Additionally, on each trial, mice move their eyes conjugately in the direction of the upcoming turn. Not only do they move their eyes, but saccadic eye movement is coordinated with the asymmetric positioning of the whiskers. Our analysis shows that the asymmetric positioning of the whiskers predicts the direction of turn that mice will make at an earlier stage than eye movement does. We conclude that, when mice move or plan to move in complex real-world environments, their motor plan and behavioral state can be read out in the movement of both their whiskers and eyes.

**Significance statement:** Natural behavior occurs in multiple sensory and motor dimensions. When we move through our environment we coordinate the movement of our body, head, eyes and limbs. Here we show that when mice navigate a maze, they move their whiskers and eyes; they position their whiskers asymmetrically, and use saccadic eye movements. The position of the eyes and whiskers predicts the direction mice will turn in. This work suggests that when mice move through their environment, they coordinate the visual-motor and somatosensory-motor systems.

## Introduction

One of the fundamental activities of the brain is to monitor and control the movement of the body. A simple everyday act like walking is associated with a sequence of movements that involve the body, limbs, head and eyes (Bizzi et al., 1971; Grasso et al., 1988; Dietz et al. 2001; Dietz, 2002; Land 2006; Foulsham, 2015). Despite the obvious importance of understanding the coordination between various sensory–motor elements engaged in planning and coordinating behaviors, the details of this coordination are still not completely clear.

In rodents, exploration of the environment is often linked to movement of the head and whiskers (Arkley et al., 2014; Towel and Hartmann, 2006; Hartmann et al., 2003; Sofroniew et al., 2014), with facial movements linked to activity observed in widespread areas of cortex (Churchland et al., 2018; Stringer et al., 2019; Steinmetz et al., 2019). Eye movements – which are thought to have “just in time” and “look-ahead” function in many species (Land and Hayhoe, 2001; Land, 2006; Mennie et al., 2007; Land, 2009; Foulsham, 2015; Srivastava et al., 2018) – have often been overlooked in rodents. Part of the reason for this neglect is related to the difficulty in measuring the motion of the small rodent eyes (Payne and Raymond, 2017; Meyer et al., 2018; Meyer et al. 2020). Additionally, rats and mice are nocturnal animals with a highly developed somatosensory system (Woolsey and Van Der Loos, 1973), where the relationship between whisking and navigation is observable more easily (Arkley et al., 2014; Towel and Hartmann, 2006; Hartmann et al., 2003; Sofroniew et al., 2014; Dominiak et al., 2019).

In the last decade there has been an evolution in our thinking as it has become increasingly obvious that rodents use their visual system to guide their response to predators and move their eyes to guide navigation (Yilmaz and Meister 2013; Wallace et al., 2013; Meyer et al., 2020). Just as in other species, rodents move their eyes when they move their head, and even though movement of the eyes is diminished when mice are head fixed, frontal cortical circuits for eye movement exist in mice; head-fixed mice can even be trained to move their eyes to target locations (Itokazu et al. 2018; Sato et al., 2020). Taken together it is likely that even in rodents, eye movement is coordinated with actions in other sensory motor dimensions, i.e. for motor planning, and for coordinating movement during navigation (Land, 2009).

In our previous work we have shown that when mice plan their movement in a real-world floating maze, they begin to position their whiskers asymmetrically. Whisking and whisker asymmetry were related to the behavioral state of the animal, and predicted turn direction that mice imposed on the maze. Mice were not explicitly trained to use their whiskers, their ability to obtain a reward did not depend on detecting tactile features of the maze (Dominiak et al., 2019). Here we have extended our earlier work by training mice to overcome their natural handedness preference. Our work shows that asymmetric whisking was related to the behavioral state of the animal — occurring in a manner consistent with the cued-direction of the upcoming turn — and was coordinated with eye movement. Mice moved their eyes and whiskers in a behaviorally-relevant, “look-ahead” fashion.

## Methods

### Behavioral experiments

We performed all procedures in accordance with protocols for the care and use of laboratory animal approved by the Charité–Universitätsmedizin Berlin and the Berlin Landesamt für Gesundheit und Soziales (LaGeSo).

#### Surgery

Adult C57bl6 mice (n=9), weighing 25 to 32 g were anesthetized with Ketamine/Xylazine (90mg/kg / 10mg/kg). Lightweight aluminium headposts were affixed to the skull using Rely X and Jet Acrylic (Ortho-Jet) black cement (Ebner et al., 2019; Dominiak et al., 2019). In the two days after surgery, analgesia was provided by Buprenorphine and Carprofen injections.

#### Airtrack

The Airtrack consists of 3 parts: 1) A square plexiglass airtable with tiny, evenly-spaced holes for holding the platform, a circular maze aloft. 2) A lightweight, circular maze that floats on the bed of air created, and 3) A pixy camera that tracks the position of the maze (Nashaat et al., 2016; Dominiak et al., 2019). Each hole of the airtable has a plastic ball bearing that moved to the mouth of the hole preventing air flow when the platform is not covering that particular hole, which reduced the hissing sound associated with the outflow of pressurized air. For the floating-maze, we used a smaller, and lighter platform than the original Airtrack plus maze which weighed 160 g and was 30 cm in diameter (Dominiak et al., 2019). Here we used a styrofoam base with a milled-out Plexiglas ring on the borders that reduced friction with the walls of the airtable. The plus maze used here had a diameter of 22 cm and weighed 30 g. It had four identical, smooth, 10 cm long lanes.

The pixy camera positioned under the platform used a color code (Nashaat et al., 2016) to track the position, the direction of movement, and speed of movement of the maze as it is guided along by the mouse. When mice entered the correct lane, the pixy output was used to trigger a motor that lowered the lick spout into position.

#### Training

One week after surgery, mice were habituated to being handled, and to the Airtrack plus maze platform. In the first days of habituation, mice acclimated to having their head post handled by the experimenter, and to short (up to several minutes) head fixation. In the course of the first week of habituation, duration of head fixation was gradually increased from 5 to 40 minutes. In addition to head fixation, mice were habituated to having their whiskers painted. During whisker painting, mice were rewarded with sweetened condensed milk.

After a week of habituation, mice were water deprived and were trained to move the platform around themselves. At the beginning of this training phase, the experimenter manually nudged or guided the animal. Over the course of 14 consecutive days of training, mice learned to move the maze by themselves, and gradually increased the number of trials they performed. Note that there were no temporal constraints on mice; the trials were self-initiated, and each behavioral epoch and each trial could be as long or as short as mice made them.

Mice were trained to use LED light cues to turn the maze in the cued direction (**Figure 1**). A trial began when mice finished with the reward from the previous trial and were still at the end of lane. One of two LEDs turned on, indicating the expected turn direction for the ensuing trial (**Figure 1A, B; Video 1** generated using real time tracking of pupils). The light was left on until mice had propelled themselves backwards, out of the lane and had begun to turn in the expected direction. When they were at the entrance to the correct adjacent lane, the LED light was switched off. Each trial was manually annotated and classified into behavioral epochs (**Figure 1C**): quietly sitting at end-of-lane, moving backward, turning, moving forward, waiting for reward, or licking.

**Figure 1.**
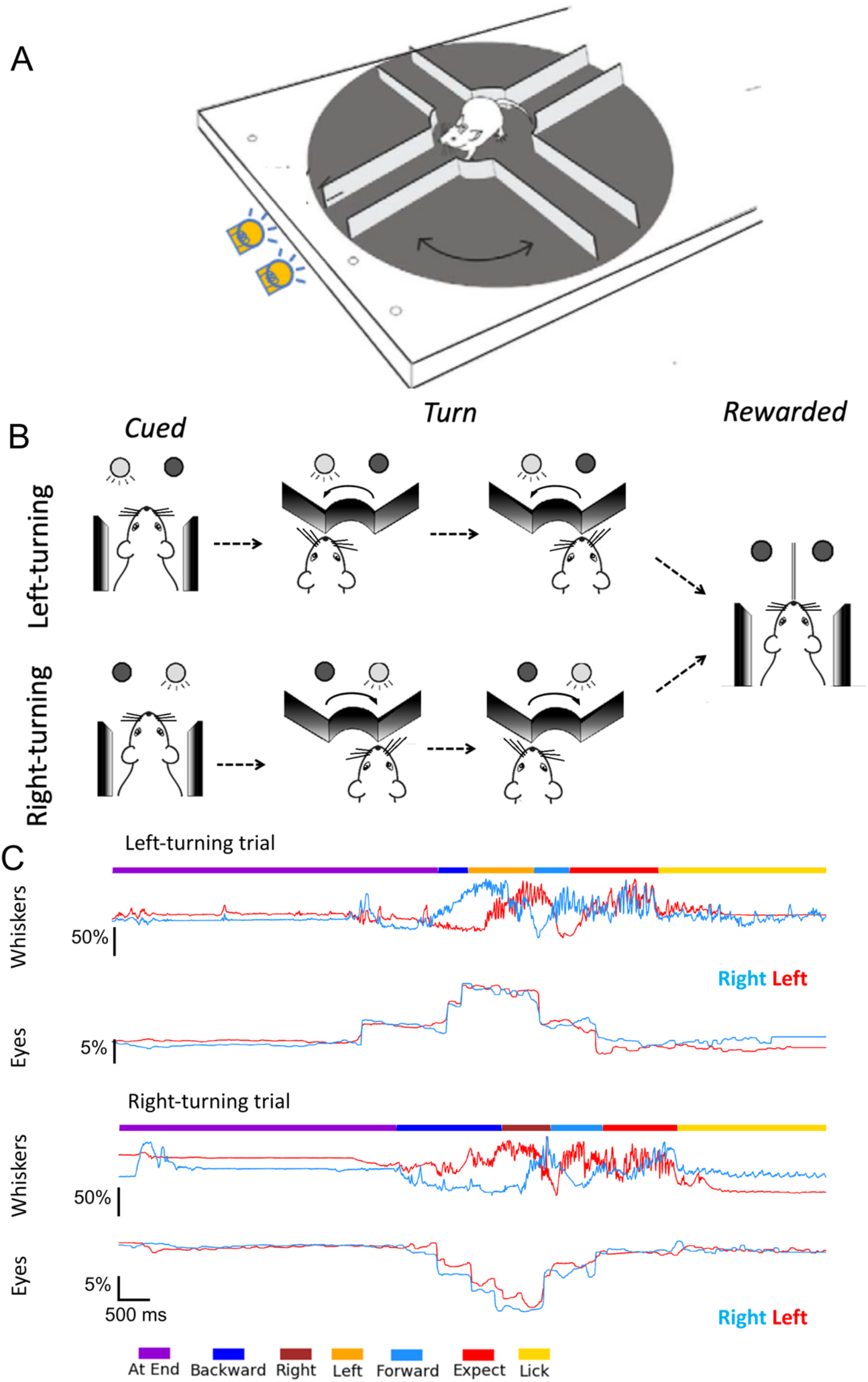
Schematic of trials, behavioral state estimates during whisker and eye movements. **A,B.** Schematic of cued directions. A trial starts with the animal at the end of the last rewarded lane. One of the two cue LEDs indicate the direction the animal should take, when they have moved backward and out of the lane. The right LED indicates that for the trial mice had to move in a clockwise direction and enter the adjacent lane where they wait for the reward tube to descend. The left LED indicated a trial with an opposite turn direction. **C.** Single right and left turn trials, showing whisker and eye movement. Mice move backward at their own pace, reach the center of the maze and turn left (top) or right (bottom) until the reward LED turns off signaling that the mouse was at the mouth of the correct, rewarded lane. Mice learn to expect a reward at the end of the lane. Once the lick spout was in their reach, mice lick the reward. During this behavior, whiskers on two sides of the face (left side whisker, blue, right side whiskers, red) move in opposite directions for left (top) and right (bottom) turns. The side to side asymmetry becomes evident just as, or just before, mice begin to move backward in the lane. By comparison, the eyes (right eye, blue trace and left eye, red trace) move in a conjugate fashion on both sides of the face. The behavioral state was manually annotated: purple, end of lane; dark blue backward movement; brown right turns; orange, left turns; light blue, forward; red, expect reward; and yellow, lick. Scale bars represent the amplitude of movement relative to the full amplitude of whisker motion (for whiskers) or relative to the width of the eye (for eyes) during the session.

#### Data acquisition

Video data from painted whiskers and the movement of the eyes were collected as mice performed the task (**Figure 1C**). While we recorded data in 3394 trials from 124 sessions in 7 animals, we manually annotated just a fraction -- 113 -- of these trials. These trials were taken from 21 sessions in 4 animals. After an additional validation step where the video data from the different streams, whisker and pupil were checked for alignment, we used 91 trials from 17 sessions recorded in 4 animals for the analysis.

To track the behavioral states and whisker positions, behavioral data was recorded at 200 Hz with a Basler acA1920-155uc USB 3.0-camera and a f=25mm / F1.4 objective, being set above the animal. The C2 whiskers on both sides were painted green (UV glow, https://www.uvglow.co.uk/), and illuminated by UV torches. Videos were acquired in the proprietary format from Matrox Imaging (https://www.matrox.com/) and later converted into the H.264 format. The acquisition and conversion were accomplished using ZR-view, a custom software (Robert Zollner, Eichenau, Germany).

For recording the animal’s pupils, two Basler acA1300-200um USB 3.0-cameras, each with an f=50mm/F2.8 objective, were used. Two independent infrared light sources (Thorlabs, https://www.thorlabs.com/; LIU850A) were directed toward the eyes. Video files of both pupils were acquired at 100 Hz using Pylon-PD, a custom software made by Eridian Systems, D-10179 Berlin, Germany.

An Atmel ATmega328P was reprogrammed to generate synchronized 100 Hz and 200 Hz trigger pulses for simultaneous frame acquisition from the three cameras (i.e. 2 pupil cameras and the behavioral camera). The Arduino Mega board that monitored Airtrack movement was also used to control the trial-based data acquisition. Trials were separated by two consecutive TTLs pulses; the first TTL marked the end of a trial and the second one marking the start of a new trial. These TTL signals were used to trigger the start and the end of video acquisition. Behavioral states in each trial were annotated manually using the behavioral video files, by marking the frames when state transitions occurred. Entry into or exit from a lane were annotated by using the position of the nose in relation to the edges of the lanes. The frame on which the animal started moving continuously in one direction was defined as the onset of forward or backward movement.

### Data analysis

#### Data selection and analysis

The 91 trials taken from 17 sessions in 4 animals, accounted for 728 annotated behavioral epochs that were used for data analysis. The following analytical procedures were performed using Python (https://www.python.org/, version 3.7.6), along with several standard modules for scientific data analysis (NumPy, https://www.numpy.org/, version 1.18.1; Scipy, https://www.scipy.org/, version 1.4.1; matplotlib, https://matplotlib.org/, 3.1.3; pandas, https://pandas.pydata.org/, 1.0.1; scikit-learn, https://scikit-learn.org/stable/, version 0.22.1; Bottleneck, https://pypi.org/project/Bottleneck/, 1.3.2) and non-standard packages (sliding1d, https://github.com/gwappa/python-sliding1d, 1.0; fitting2d, https://doi.org/10.5281/zenodo.3782790, 1.0.0a2).

#### Tracking whisker movement

Whisker positions were estimated using the custom Python script (videobatch) (DOI:10.5281/zenodo.3407666). The top-view videos first underwent a maximum intensity projection using the videobatch script. Regions of interest (ROI) for tracking were selected manually for the C2 whiskers on both sides of the face using Fiji’s freehand selection tool (Dominiak et al., 2019; Sehara et al., 2019). Using the Python script, pixels that belonged to a particular hue value were collected and the luma-weighted average position was computed. For frames where the algorithm failed for any reason, values were dropped and were filled in later by linear interpolation.

The whisker positions tracked during each behavioral session were then converted to whisking angles. Using the fitting2d Python library, a circle was fit to the set of two-dimensional positions for each whisker, and the position at each timepoint was converted into the polar coordinates around the fitted circle. The angle corresponding to the session-median position was defined as the zero angle for each session.

#### Tracking eye movements

DeepLabCut (DLC; version 2.1) (Mathis et al., 2018; Nath, Mathis et al., 2019) was used to track pupil positions from video frames. A deep-neural network model was trained to detect eight points on the edge of the pupil on each video frame. A single model was used for tracking pupils of both sides of the face of all animals in all sessions. In total, 284 frames from 28 sessions from different animals were manually annotated to train the network. For each video frame, the eight edge points of the eye detected by the deep-neural network were then fitted by an ellipse, using the fitting2d Python library. The position and the diameter of the fitted ellipse were considered to be those of the pupil.

Because the size of the eyes in the field of view of the video varied across behavioral sessions, the position of the pupil was normalized with respect to the size of the eye. For each eye during each behavioral session, the average eye shape was first delineated manually using the ROI Manager of ImageJ, based on the average-projection image of eye videos during the session. A parabola curve was then fitted to the trace corresponding to either the top or the bottom eyelid, using the fitting2d Python library. The two corners of the eye were then computed as the crossing points between the two fitted parabola curves. These defined the line segment representing the width of the eye. The position of the pupils was first projected onto the eye-width segment, and then represented in terms of the fraction relative to the full width of the eye (hereafter called as normalized pupil position).

#### Detection of saccades

The first derivative (dX / dt) of normalized eye positions was computed to obtain the pupil speed. Sudden changes in eye position appeared as distinct spikes in the corresponding eye speed trace. Saccades were said to have occurred when the absolute values of eye speed for both eyes crossed a threshold value which was set empirically at 0.1% (relative to the full eye width) per frame. If a train of these high speed events occurred within an interval of less than 250 ms, it was considered to be a processing artifact and was discarded from the analysis.

#### Normalization of behavioral state durations

The duration for each behavioral state was normalized using interpolation. For each behavioral epoch, we first set up a normalized time base such that the timepoints 0 and 1 marked the start and the end of the epoch. The data points were resampled from the original time base (i.e. frames) to the normalized time base using interpolation.

#### Analysis using ROC curves

We used ROC curves to examine whether eye movement or whisker asymmetry or both could predict the turn direction. We divided the backward-movement behavioral epoch into 5 distinct time bins. For each time bin, we set up a set of classifier models. As its input, each classifier model received either a whisker asymmetry value (i.e. the right-whisker protraction subtracted by the left-whisker protraction) or an eye position value (positive values indicating leftward deviations and negative values indicating rightward deviation). Each model had a certain pre-specified threshold value. When input value was above the threshold value, the model determined that a left turn would occur. If the input value was below the threshold value the model predicted a right turn. We supplied the set of models of different threshold values with behavioral data (either whisker asymmetry or eye positions). For each time bin, the probability of each model’s correct prediction of the upcoming turn direction to the left, the “true positives”, and the probability of predicting the upcoming turn direction incorrectly, the “false positives” (the model predicted to turn left, whereas the animal turned right in reality) were computed. The ROC curve for each time bin was then plotted based on these probabilities, using the set of models having different threshold values. The area-under-curve values generated by this analysis were compared to the shuffled data, and significance was assessed for each time bin, for whisker asymmetry and eye movement separately. Statistical comparison between the raw and shuffled data and between whisker-based and eye-based data in each time bin was performed using Mann–Whitney U-test. Mann– Whitney U-test with Bonferroni correction was used for pairwise comparison between data belonging to different time bins.

## Results

### Whisking asymmetry and eye position

To examine whether the positioning of whiskers asymmetrically reflected the upcoming turn direction — whether whisker asymmetry could be reset from trial to trial — we tracked whisker movement bilaterally as mice performed the task. To examine the relationship between eye movement in locomotion and task performance, we also tracked eye movement bilaterally.

As we have shown earlier, mice whisked when they were engaged in active behaviors, and they did not whisk much when they were inactive (**Figure 1C,** Dominiak et al., 2019). Whisking continued as long as mice were moving, and decreased when mice were expecting reward and licking. But whisking was not uniform bilaterally; mice whisked asymmetrically, they moved their right and left whiskers in opposite directions throughout the course of the trial almost as long as they were moving in the maze (**Figure 1C** top, a left turn trial, and bottom a right turn trial). As mice backed out of a lane, turned and went forward into a new lane (**Figure 1C**, *orange* for left turn, and *brown* for right), the asymmetry flipped. Asymmetry of whiskers reflected the direction of turn (**Figure 1C***top* left turning trial, and *bottom* right turning trial), with the right and left side C2 whiskers mirroring each other for the right and left turns. These single trial data from a mouse, moving to the right or left on some trials, suggest that whisking asymmetry arose as part of the motor plan.

In addition to positioning the whiskers asymmetrically and whisking on every trial, mice also moved their eyes on every trial (**Figure 1C; Video 1**). They moved their eyes rapidly when they were moving, i.e. when they were engaged in active behavior. The movement / position traces of both eyes were similar, seen by the overlaid traces for each eye (red and blue traces, **Figure 1C**); mice moved their eyes conjugately, bilaterally, to the same extent, at roughly the same time, in the direction of the upcoming turn. Consequently, right and left turn trials elicited movement of both eyes, but movement was in the opposite direction for the two turn directions. Once mice stopped moving — when they were expecting a reward, licking, or were waiting to begin the new trial — movement of the eyes diminished greatly.

These single trial data suggest that both whisker and eye movements were dependent on the behavioral state of the animal, i.e. whether they were active and moving or quietly sitting. To quantify these effects, we normalized the data for time (each trial, and each behavioral epoch could have different duration) and normalized whisker position and eye position data for extent / amplitude of movement (**Figure 2**). These normalized average traces confirm what the single trial data show: the side-to-side whisker asymmetry (**Figure 2A**) and eye movement (**Figure 2B**) were related to the animal’s behavioral state, and direction of turn in the maze. When right-turn trials were compared to left-turn trials, whisker asymmetry appeared as a mirror image and eye movement traces appeared as inverted mirror images of each other. Whiskers on two sides of the face moved bilaterally but in the opposite direction, and at the same time mice moved their eyes conjugately in the same direction. This pattern of whisking and eye movement were evident when mice were in an active state, i.e. when the animals were moving backward, forward or turning and the pattern was not evident when mice were standing still at end of lane, expecting reward or licking. When mice were active and moving backward, forward or turning, in both right and left turning trials, there was significant whisker asymmetry and significant change in eye position compared to behavioral epochs where mice were just standing still, expecting reward or licking (p < 0.05, Wilcoxon sign rank test). Taken together these data indicate that whisker asymmetry and the movement of the eyes were active processes occurring in a behaviorally relevant manner as mice navigated the plus-maze.

**Figure 2.**
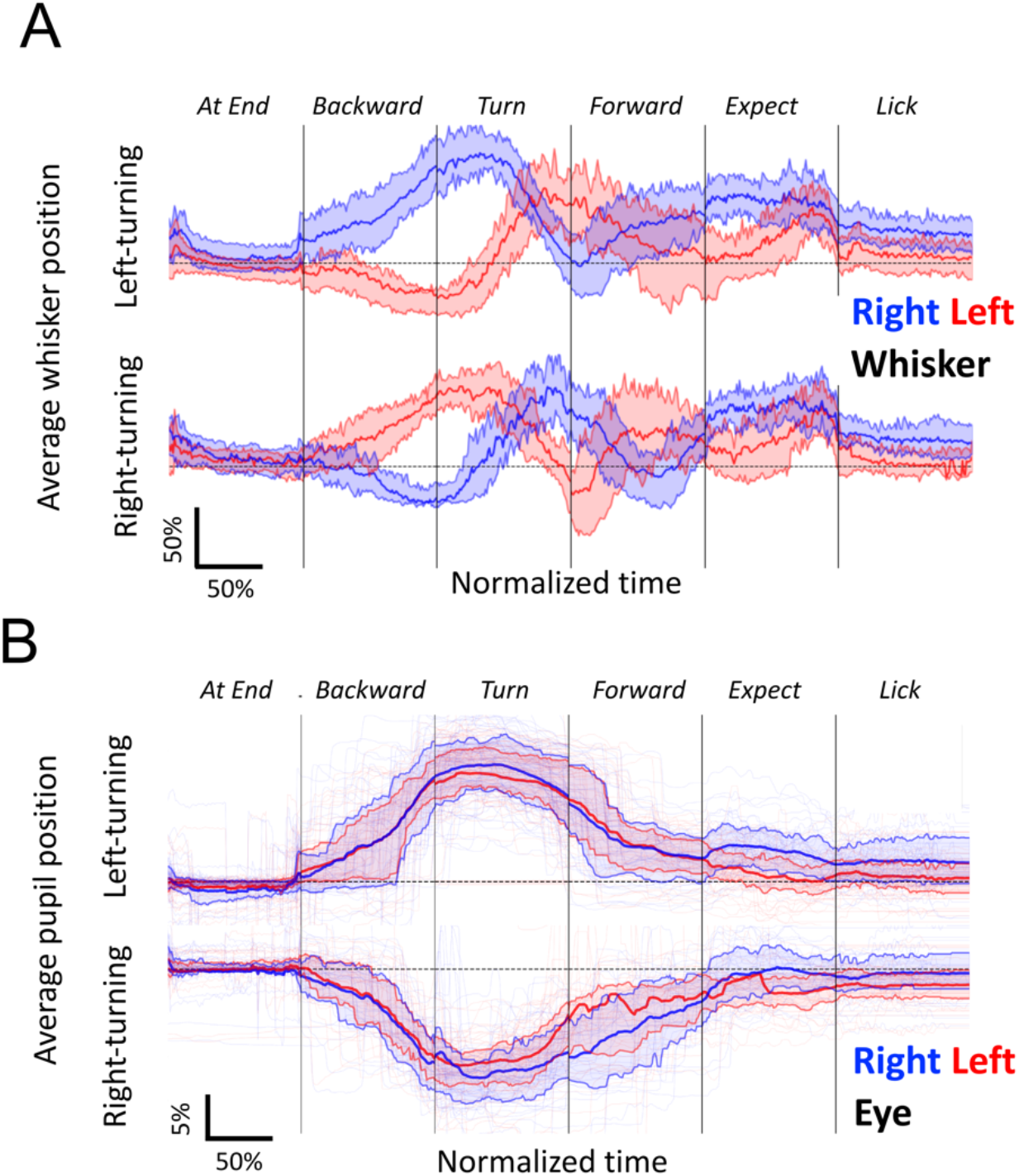
Whisker asymmetry and eye position to the cued direction of movement. **A.** Averages of whisker positions. Median positions for the left whisker (thick red line) and right whisker (thick blue line) show that whisker asymmetry was related to direction of movement, and was evident early in the backward movement of mice out of the lane. The shaded areas and thinner lines show the 25 to 75 percentile. The asymmetry between whiskers emerges at the beginning of backward movement, reverses once during the course of the turn, a second time during the forward motion. Asymmetry vanishes when mice wait for and expect reward. Note that whisker asymmetry on right- and left-turn trials are almost mirror images of each other. **B,** Averages of eye positions. Median positions for the left eye (thick red line) and the right eye (thick blue line) move conjugately in the behavioral epoch-specific manner. Note that the average eye positions on right- and left-turn trials are inverted images of each other. Whisker and eye positions were normalized for amplitude (see Methods) and for time (because trial durations varied, see Methods). The numbers of behavioral epochs used were: AtEnd, n=54 (left-turning) and 36 (right-turning) epochs; Backward, n=54 (left-turning) and 36 (right-turning) epochs; Turn, n=81 (left-turning) and 58 (right-turning) epochs; Forward, n=51 (left-turning) and 38 (right-turning) epochs; Expect, n=51 (left-turning) and 38 (right-turning) epochs; Lick, n=51 (left-turning) and 38 (right-turning) epochs.

### Predicting turn direction with whisking and eye movements

We used ROC curves to examine whether eye movement or whisker asymmetry or both could predict the turn direction (**Figure 3**, also see Materials and Methods). When normalized for time (during the backward movement epoch) and binned, the ROC curves reveal that both the whisker asymmetry and eye position become increasingly stereotyped and increasingly predict the turn direction (**Figure 3A**).

**Figure 3.**
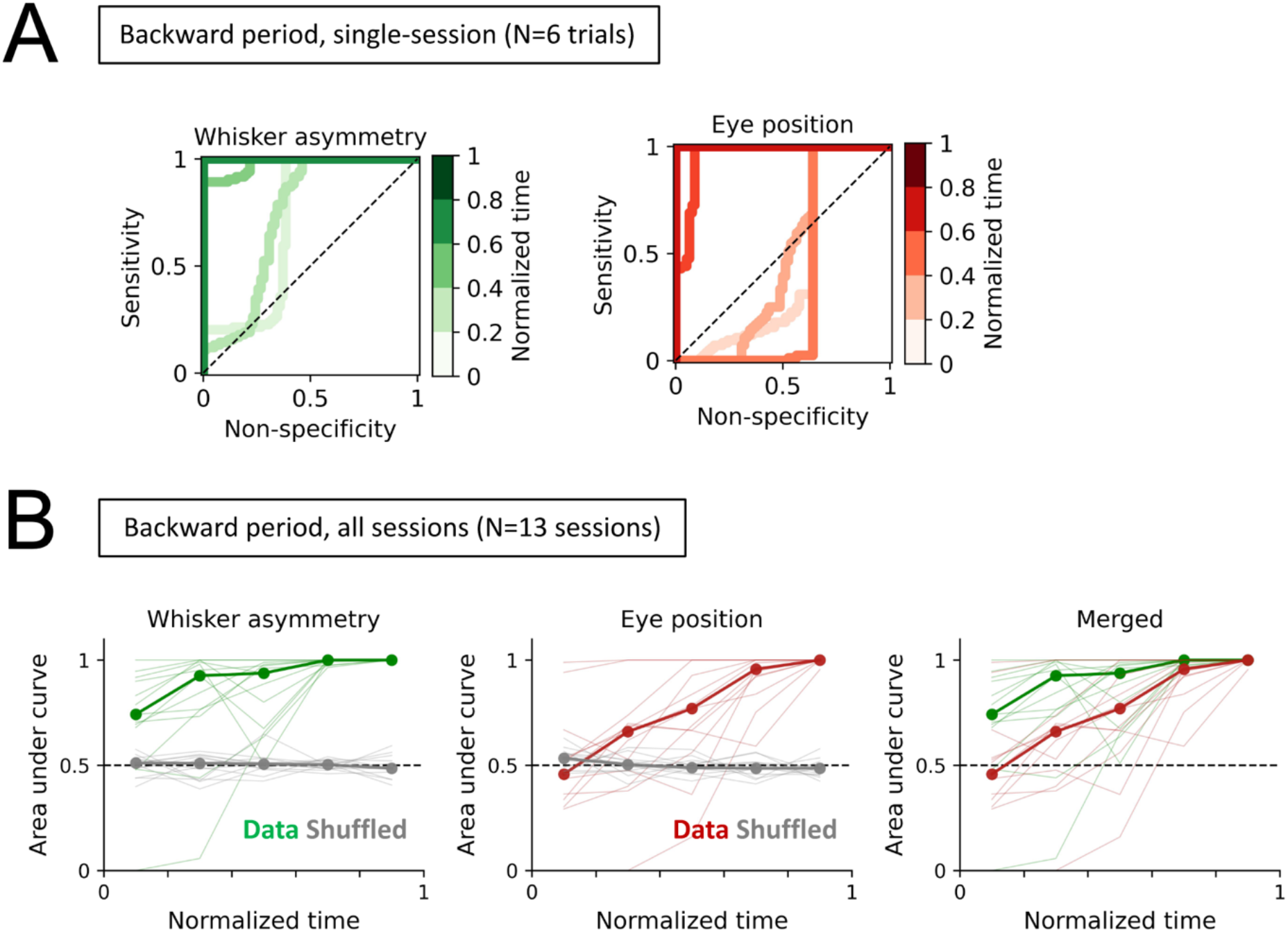
Behavior during the backward motion predicts the next turn direction of the animal. **A.** ROC curves were generated based on the whisker asymmetry (left, green) and the eye position (right, red) during the Backward period to predict whether the animal was going to turn in the left or right. The denser curves represent the later phases in the backward movement epoch. Note that during the later phases, the curves are positioned closer to the top-left corner of the bounding square indicating that whisker asymmetry and the eye position are good predictors of the upcoming turn direction. Data from a representative session are shown. **B.** The area under curve (AUC) at different phases during the backward movement epoch in multiple behavioral sessions and animals (n=13 sessions taken from 4 animals, consisting of 77 Backward epochs in total) show that the whisker positions and eye positions separately work as an efficient predictor of the animal’s upcoming turn direction. The median AUC values were computed for whisker asymmetry (left, thick green line) or for the eye position (center, thick brown line). The gray lines on the two left panels represent the AUC values when the whisker asymmetry or the eye position data were randomized within the backward period. The right panel shows the comparison of predictive performances between the whisker asymmetry and the eye position on different phases within the backward movement. Pairwise comparisons indicate that whisker asymmetry-based prediction was significantly better for later time bins than for earlier ones, reaching significance midway through backward motion: bin #1 vs bin #2, p=0.7190; #1 vs #3, p=0.5040; bin #1 vs bin #4, p=0.0067**; bin #1 vs bin #5, p=0.0002***; bin #2 vs bin #3, p=1.0000; bin #2 vs bin #4, p=0.1119; bin #2 vs bin #5, p=0.0030**; bin #3 vs bin #4, p=0.0971; bin #3 vs bin #5, p=0.0062**; bin #4 vs bin #5, p=0.0263*; Mann–Whitney U-test with Bonferroni correction. Pairwise comparisons for eye movement also indicate that late in the backward motion eye movement was a good predictor of upcoming turn direction: bin #1 vs #2, p=1.0000; bin #1 vs #3, p=0.9991; bin #1 vs #4, p=0.0147*; bin #1 vs #5, p=0.0001***; bin #2 vs #3, p=0.8704; bin #2 vs #4, p=0.0689; bin #2 vs #5, p=0.0046**; bin #3 vs #4, p=0.5576; bin #3 vs #5, p=0.0085**; bin #4 vs #5, p=0.0671).

While both eye movement and whisker movement accurately predicted turn direction before the animal started to turn the maze, examination of the area under curve (AUC) across behavioral sessions (n=13 sessions taken from 4 animals, consisting of 77 Backward epochs in total) showed that the AUC for models based on whisker asymmetry was significantly larger than the chance level from the very beginning of the backward-movement period. This implied that whisker asymmetry was a good predictor of the upcoming turn direction (bin #1, p=0.0024; bin #2, p=0.0015**; bin #3, p=0.0001***; bin #4, p=0.0000***; bin #5, p=0.0000***; Mann– Whitney U-test, **Figure 3B, left**). Pairwise comparisons indicate that whisker asymmetry-based prediction was significantly better for later time bins than for earlier ones, reaching significance (p< 0.01, Mann–Whitney U-test with Bonferroni correction) midway through the backward motion (**Figure 3B, left**).

Eye position also correctly predicted turn direction. Except for the beginning of the backward movement, eye position also predicted the upcoming turn direction (bin #1, p=0.3222; #2, p=0.0406*; #3, p=0.0015**; #4, p=0.0000***; #5, p=0.0000; Mann–Whitney U-test). As the animal moved closer to the turn, during or just after the mid-point of the backward motion, eye position became a good predictor of the upcoming turn direction (p < 0.05, Mann Whitney U-test with Bonferroni correction, **Figure 3B, center**).

Additionally, models based on whisker asymmetry were slightly but significantly better at predicting turn direction — just as the animal starts the backward movement — than those using eye positions (bin #1, p=0.0146*; bin #2, p=0.0153*; bin #3, 0.1772; bin #4, 0.0353*; bin #5, 0.1780; Mann–Whitney U-test) (**Figure 3B, right**). These analyses reveal that the information in whisker asymmetry and eye position, individually and together were sufficient to predict the turn direction before the animal begins to turn.

### Saccadic eye movement and behavioral state

While the eye movement data from single trials shows that head-fixed mice moved their eyes in a stepwise fashion on each trial (**Figure 1**), this fast movement of the eyes tended to be smoothed out in the average traces (**Figure 2**). To examine the relationship between the rapid saccadic eye movements and behavioral state, we related saccades which appeared as distinct spikes in the corresponding eye speed trace to the behavioral state (**Figure 4A**). Eye movements were counted as saccades if the following criteria were met: absolute value of eye speed for both eyes was above the threshold of an absolute eye speed of 0.1% of full eye-width per frame, and there was no additional spike of higher absolute speed related to eye movement within 250 ms of the saccade. The duration of each behavioral epoch (At End, Backward, Turn and Forward) was normalized and divided into 10 equal bins, and the number of saccades that occurred within each time bin were counted. This analysis revealed that mice made saccades predominantly in the latter half of the backward movement, as they moved out of the lane. Saccades also occurred as mice turned and went forward into an adjacent lane (**Figure 4B**). Consistent with the analysis of eye positions above in **Figure 1C** and **Figure 2**, saccades were more likely to occur in the direction of the upcoming turn, rather than in the opposite direction. The distribution of saccades suggests that mice look in the direction of turn as they exit backward out of the lane, and again as they enter another lane. Note that although saccades occurred primarily when the mice were close to a turn or turning saccades also occurred when the animals were merely moving forward or backward in a lane.

**Figure 4.**
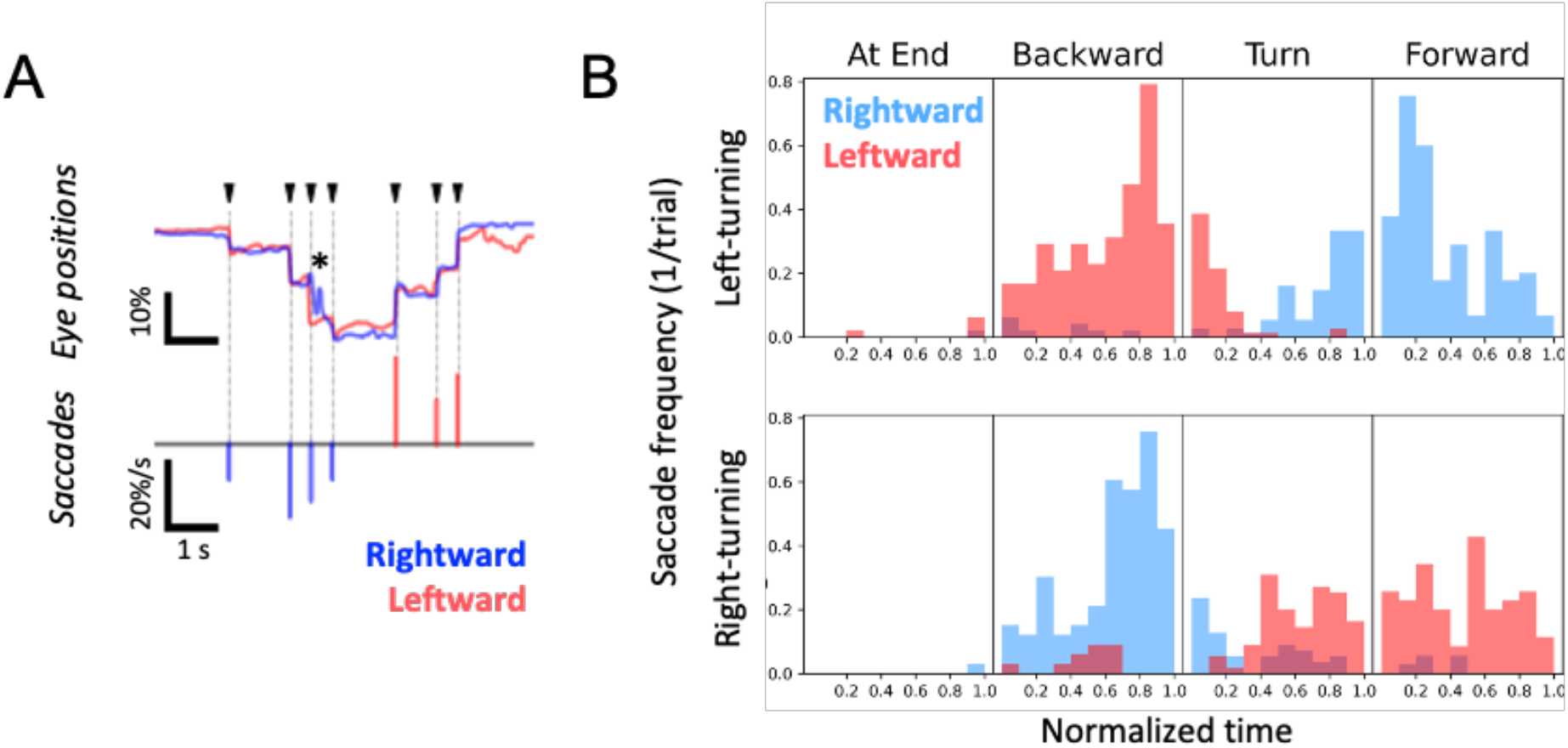
Saccadic eye movements related to behavioral state and turn direction. **A.** Detection of saccades. Saccadic events were defined as the time points when the two eyes (red and blue traces) moved rapidly and conjugately (arrowheads). Asterisk refers to an occlusion-related artifact where one eye was partially occluded by the movement of a wall. **B.** Timings of saccades were related to the behavioral state, and saccade direction was related to turn direction. Histograms of per-epoch saccade occurrences were generated based on the data of 91 trials from 17 sessions taken from 4 animals. Leftward saccades (red bins) occurred as the mice turned leftward (top distribution), when they backed out of a lane, and rightward saccades (blue bins) occurred when they turned right and went forward into a lane. The direction of saccades was inverted for right turning trials (bottom distribution). Saccades were detected mainly when animals were moving and active. The timings of saccades distributed broadly across different phases during each behavioral epoch.

### Correlation between saccades and whisker asymmetry

In many species, eye movement is correlated with head movement (Land, 2018). In rodents, head movement has often been related to both eye and whisker movement (Meyer et al., 2019; Hartmann et al., 2003; Towel and Hartmann, 2006). But mice can move their whiskers and eyes independently of their heads, even when they are head fixed, so it was possible that even though movement of the head was prevented, eye and whisker movements could be coordinated. To examine the relationship between saccades and whisker movement, we plotted the level of asymmetry in the 500 ms before and after saccadic eye movements as mice began to move backward out of the lane (**Figure 5**). We used the beginning of the backward motion epoch because it elicited a number of saccades, providing a decent sample size. Additionally, in this phase of the trial, mice mostly avoided contact with the walls of the maze, which also meant that we had a mostly unobstructed view of both whisker positions and eye positions in the course of the backward movement.

**Figure 5.**
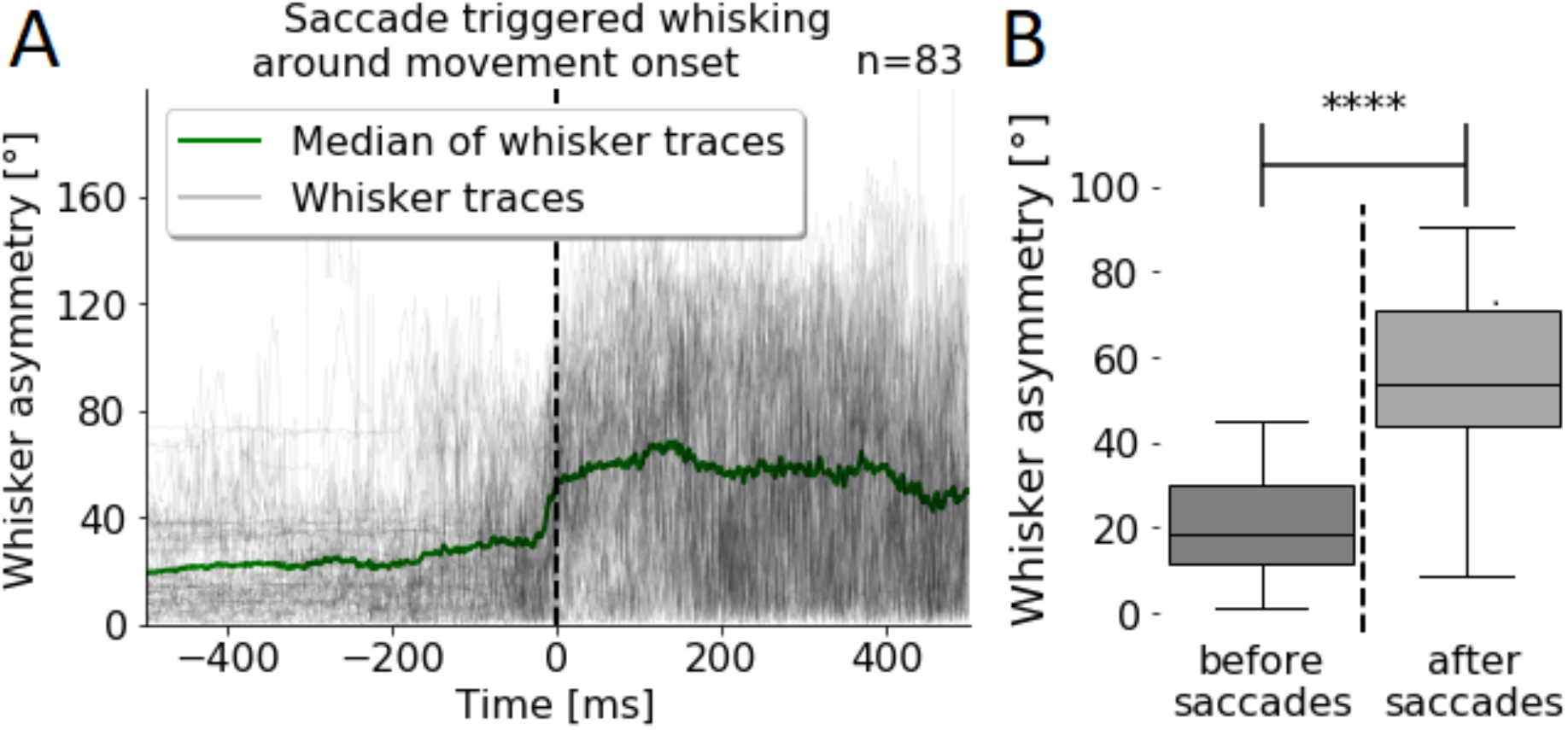
Coordination between saccadic movement and whisker asymmetry. **A.** Changes in whisker asymmetry in relation to the occurrence of saccades around the onset of movement at the end of the lane. The dashed line at zero marks saccade onset. On average, whisker asymmetry increased just around the time the eyes moved (n=83 saccade events). The increased asymmetry was maintained for hundreds of milliseconds after each saccade. Individual traces of whisker asymmetry related to saccade onset are plotted in black, and the median is shown in green. **B.** Quantification of changes in whisker asymmetry. The difference in whisker position, measured in degrees increases significantly after saccades occurred (Wilcoxon signed rank test, p < 0.0001).

Our analysis revealed that as mice moved backward and moved their eyes into position for the upcoming turn, they also changed how they positioned their whiskers. Whisker asymmetry increased measurably, almost tripling from 18 degrees to 53 degrees (**Figure 5A-B**, Wilcoxon signed-rank test, p<0.001) just after mice made saccades during their backward movement. These results suggest that mice coordinate eye movement with asymmetric positioning of the whiskers as part of the motor plan and execution of the movement related to the turn.

## Discussion

To behave means to plan and emit a sequence of actions (Tinbergen, 1955; Krakauer et al 2017). On one hand, the behavior we used here was complex; mice were head fixed as they moved their entire body through a “real-world” maze, as they pushed and rotated the platform around themselves, and entered or exited lanes. On the other hand, the behavior was simple; mice were in an environment that they controlled, it had lanes, similar to the burrows that mice tend to live in their natural habitat. All mice had to do in the task was to keep track of a cue that indicated the direction to move in; then they had to recognize an adjacent lane and enter it to obtain a reward. There was no time pressure, no requirement for mice to discriminate one thing from another. The goal was just to forage for food and water. In the course of this behavior, as part of their motor plan, mice moved their whiskers and eyes in anticipation, in preparation for a turn in the maze. The bilateral positioning of the whiskers and eyes predicted the turn direction mice expected to impose on the maze. Once the eyes moved, whiskers were positioned even more asymmetrically (**Figure 3, 6**).

**Figure 6.**
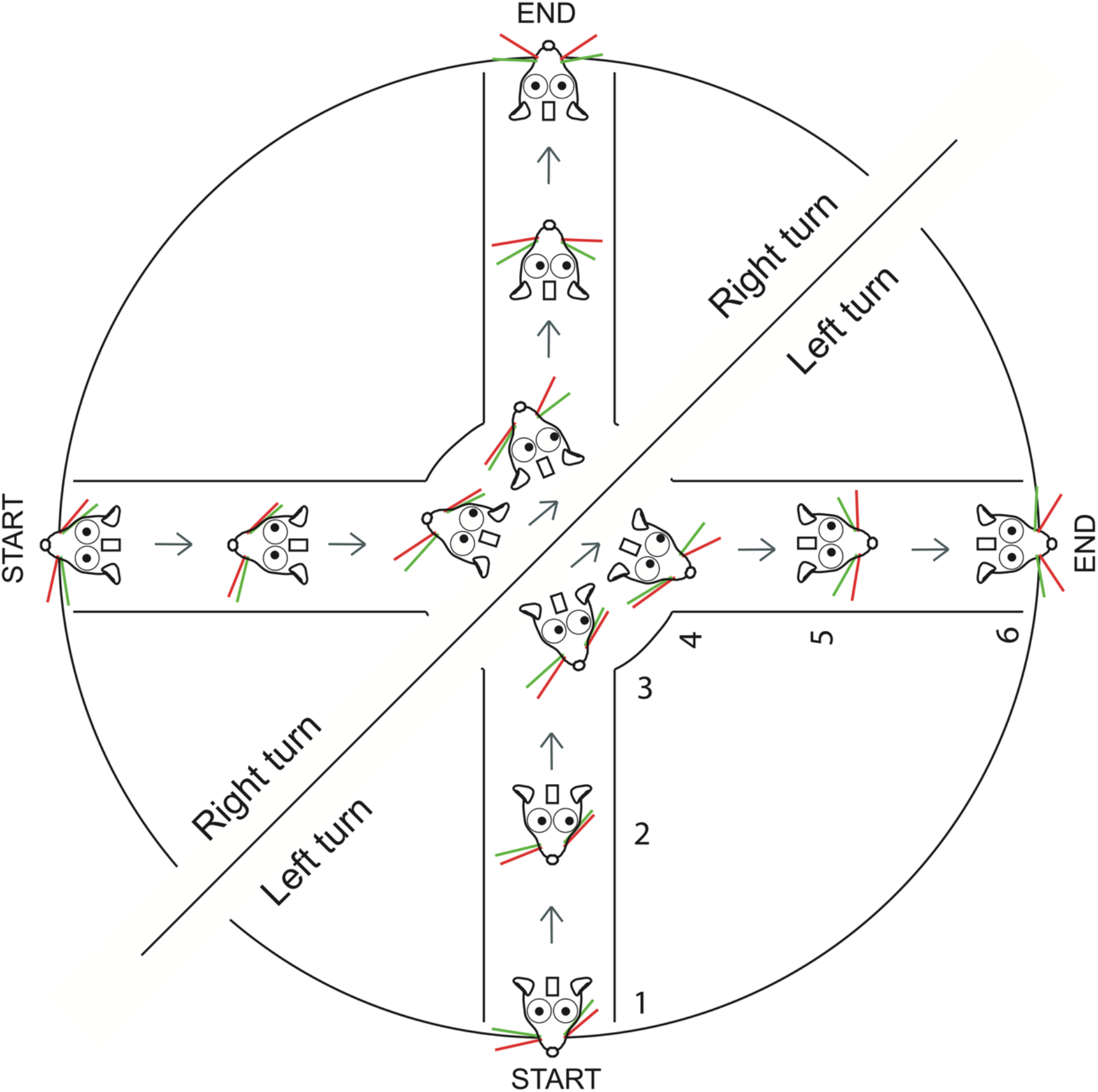
Schematic of eye movement and whisker asymmetry in the course of right and left turns. Whisker asymmetry and the eye movement in the course of single right or left turns.

In this maze — a maze with real, not virtual walls — mice moved their body, and while they moved their body, they also moved their eyes and whiskers bilaterally. In fact, our earlier work showed that in the course of this behavior mice, even move their nose and reset the set point of their whiskers, presumably by moving their whisker pad (Dominiak et al., 2019). One key difference between the earlier work and this work is that, in the earlier work, we made no attempt to condition mice to attend to cues, or to move in one of two directions based on a cue. In the earlier work, mice mostly moved in their preferred direction, right or left, but not in both directions. Nevertheless, our work suggests that in the course of this behavior mice move their whiskers, whisker pad and eyes bilaterally. They probably move their entire face in a coordinated fashion, related to their behavioral state, i.e. whether the mouse was standing still, or moving backwards etc.

The purpose of this coordination is not completely clear. Our work suggests that eye movement could have a look-ahead function for assessing the distance to a wall or to an opening. Whisker movement could also have a look-ahead function: whiskers could be positioned to anticipate the upcoming turn. But whether mice innately coordinate these movements, or they first learn to coordinate the movement of eyes, whiskers / face when they learn to navigate the maze is not clear. Whether mice coordinate whisker and eye motion in their natural, freely behaving condition is also not known, in part because the entire behavioral repertoire that we have measured here is not easily or reliably measured in freely moving animals. We also do not know whether circuits for eye movement, whisker movement, motor planning, and body movement are all active simultaneously in adjacent parts of anterior frontal and sensorimotor cortices, and how activity in these circuits interacts with subcortical circuits to generate the sequence of movements.

### Eye movement in mice

Rodents use vision to locate objects and to avoid airborne (Morris et al., 1979) or ground-dwelling predators (Doncanster et al., 1990). Freely behaving mice move their eyes when they move their head; they move their head and eyes to monitor looming stimuli, and to coordinate freezing or escaping behaviors. It has also been shown that freely moving rodents move their eyes in a conjugate or dis-conjugate fashion (Sakatani and Isa, 2007; Wallace et al., 2013; Wang et al., 2015, Payne and Raymond, 2017; Samonds et al., 2018; Meyer et al., 2018). The small, slow dis-conjugate movements of the eyes occur more frequently in freely moving rats and mice. When rodents are head-fixed, the frequency of eye movement is reduced and eye movement becomes almost completely conjugate (Wallace et al., 2013; Samonds et al., 2018).

Our work here confirms the earlier work, showing that head-fixed mice do indeed move their eyes, and they move their eyes conjugately. We extend the earlier work in two ways; first, we show that even though mice are not instructed to move their eyes (Itokazu et al., 2018; Sato et al., 2019), mice move their eyes reliably, in a behaviorally relevant fashion just before and during turns. Mice moved their eyes in anticipation of the turn, in the direction of the turn on every trial. Secondly, we show that eye movement was embedded in the coordinated motion of the animal. When combined with our earlier work, this work shows that when mice move, they move their eyes, whiskers, whisker pad and nose. In this floating real-world maze, movement of the eyes was part and parcel of the concerted movement of the face and body. Currently, we do not know whether mice move their eyes in a similar fashion in virtual reality systems that have mazes, or virtual reality systems that have visual streaming built into them, but this should be an avenue of future investigations.

### Sensory–motor coordination in the real-world environment

Simultaneous and continuous coordination of movement and sensation is part of the natural function of the brain (Welker, 1964; Llinas, 2001; Wolpert and Ghahramani, 2007; Wolpert and Landy, 2012; Musall et al., 2019; Stringer et al., 2019; Tantirigama et al., 2020). In their natural state, when animals move, they interact with their environment in multiple sensory–motor modalities; their limbs touch the floor, they look in the direction of the motion, their breathing changes and in the case of rodents, they move their whiskers. The movement of the animal changes the sensory scene for the animal, which in turn generates a new set of sensory stimuli, i.e. pressure on limbs, the novel texture under their skin, an updated stream of visual input, and tactile input from the whiskers guiding mice around obstacles around and in front of them. These changing set of stimuli elicit activity in the brain and are then used to reinforce the ongoing behavior or to modify the animal’s movement (Arkley et al., 2014; Kurnikova et al., 2017). As animals learn they can plan ahead, and anticipate what to expect before the actual sensory input occurs (Keller and Mrsic-Flogel, 2018).

Our work sheds light on the innate strategy used by mice, and shows how they spontaneously, almost automatically plan and coordinate the movement of their eyes, whiskers and body. The somewhat natural environment we used here is likely to engage widespread cortical and subcortical circuits in a manner that is close to what the brain has evolved to do, that is to move the animal through the world.

## Acknowledgements

We also thank Mostafa Nashaat and Hatem Oraby for their help at various stages of this work. We thank the Charité Workshop for technical assistance especially Alexander Schill, Jan-Erik Ode and Daniel Deblitz. Finally we thank members of the Larkum lab for useful discussions about earlier versions of this manuscript.

---

**Video 1.**
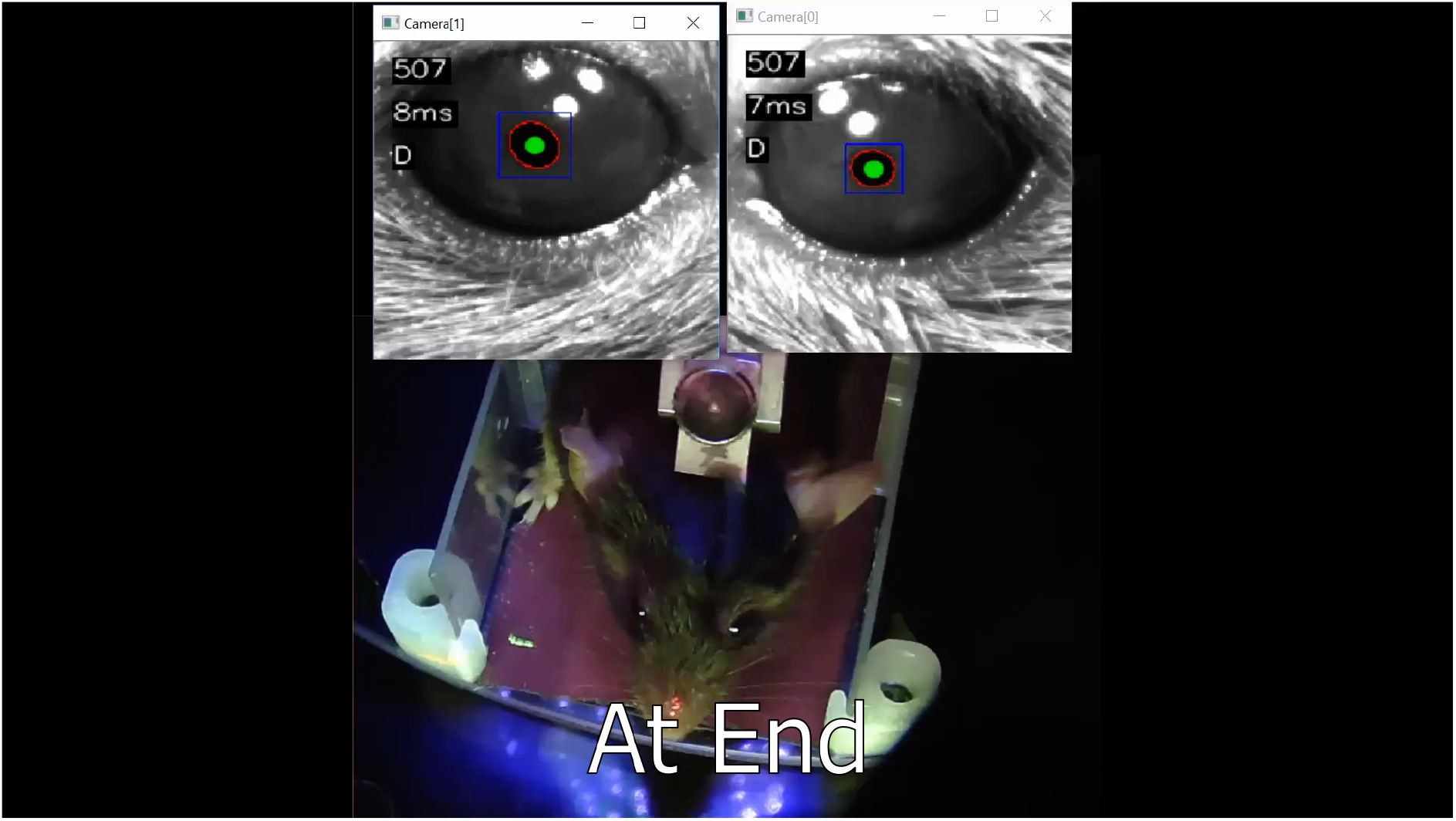
Real time eye tracking and behavior on two trials. A machine learning algorithm was used to track eye position in real-time, and to illustrate how eye position changed with the position of the mouse in the maze.

